# MetaRNN: Differentiating Rare Pathogenic and Rare Benign Missense SNVs and InDels Using Deep Learning

**DOI:** 10.1101/2021.04.09.438706

**Authors:** Chang Li, Degui Zhi, Kai Wang, Xiaoming Liu

**Author notes:** Correspondence to Xiaoming Liu, Ph.D., USF Genomics & College of Public Health, University of South Florida, 3720 Spectrum Boulevard, Suite 304, Tampa, FL 33612, USA. E-mail: C.L.; D.Z.; K.W.; X.L.

## Abstract

With advances in high-throughput DNA sequencing, numerous genetic variants have been discovered in the human genome. One challenge we face is interpreting these variants to help in disease screening, diagnosis, and treatment. While multiple computational approaches have been proposed to improve our understanding of genetic variants, their ability to identify rare pathogenic variants from rare benign ones is still lacking. Using context annotations and deep learning methods, we present pathogenicity prediction models, MetaRNN and MetaRNN-indel, to help identify and prioritize rare non-synonymous single nucleotide variants (nsSNVs) and non-frameshift insertion/deletions (nfINDELs). A recurrent neural network incorporating a +/- 1 codon window around the affected codon was combined with 28 high-level annotation scores and allele frequency features to develop the two proposed models. We use independent test datasets to demonstrate that these new models outperform state-of-the-art competitors and achieve a more interpretable score distribution. Importantly, prediction scores from the nsSNV-based and the nfINDEL-based models are comparable, enabling easy adoption of integrated genotype-phenotype association analysis methods. In addition, we provide pre-computed MetaRNN scores for all possible human nsSNVs and a Linux executable file for a fast one-stop annotation of nsSNVs and nfINDELs. All the resources are available at http://www.liulab.science/MetaRNN.

## Introduction

Next-generation sequencing (NGS) has dramatically improved our ability to detect genetic mutations in the human genome. However, our current ability to detect genetic mutations far exceeds our ability to interpret them, which is one of the significant gaps in effectively utilizing NGS data^1^. This issue is prominent for rare genetic mutations (allele frequency <1%) since traditional methods, such as population-based genome-wide association and whole exome sequencing studies, lack the power to identify the rare pathogenic or causal mutations from rare benign ones. The issue is especially prominent when the phenotype of interest has low prevalence, such as rare Mendelian disorders where only the proband’s and the parents’ genetic testing data are available. Amino acid changing mutation is probably the most well-studied candidate variant type for pathogenic mutations. However, as each healthy individual contains around 10,000 such mutations, hundreds of them are singletons^2^, it is still challenging to correctly identify pathogenic causal mutations from benign and non-functional ones in the coding regions. While being exempt from definite and severe consequences, such as mutations result in premature stop codons, non-synonymous single nucleotide mutations (nsSNVs) and non-frameshift insertion/deletions (nfINDELs) are mutations with a broad spectrum of functional consequences, which makes them even more challenging to classify.

Because experimentally validating the effects of these mutations is highly time-consuming and costly, computational approaches have been developed for this purpose^3-18^. These methods can be loosely categorized into three groups: functional prediction methods, which model functional importance of the variants; conservation-based methods, which use evolutionary data to identify functional regions and variants; and ensemble methods which combine multiple individual prediction tools into a single more powerful predictor. While these methods have been widely used to predict potentially pathogenic mutations, there are still two significant limitations in their application to whole-exome sequencing studies. First, most of these methods either deployed models trained with rare pathogenic and common benign variants or ignored the importance of observed allele frequencies as features, leading to less optimized performance for separating rare pathogenic and rare benign variants. Second, most methods provide prediction scores for only nsSNVs or incomparable scores for nsSNVs and nfINDELs separately, making it infeasible to use these scores as weights in an integrated (nsSNV+nfINDELs) burden test for genotype-phenotype association analysis.

This study developed the MetaRNN and MetaRNN-indel models to overcome these limitations, enabling users to annotate and score both nsSNVs and nfINDELs easily. As predictive features, our classifiers combine recently developed independent prediction algorithms, conservation scores, and allele frequency information from the 1000 Genomes Project (1000GP)^19^, ExAC^20^, and gnomAD^21^. Annotations from flanking +/- 1 codon of nucleotides around the target mutation were extracted by stacked bidirectional gated recurrent units^22^ (GRUs). We trained our recurrent neural network (RNN) model with 26,517 rare nsSNVs (absent from at least one of the three population datasets) and 2,057 rare nfINDELs from ClinVar^23^ up to release 20190102. To evaluate the performance of the proposed models, we compared multiple state-of-the-art computational methods using independent test datasets constructed from well-known variation-disease association databases, i.e., ClinVar^23^ and HGMD^24^, a TP53 functional mutation dataset^25^, and a dataset of potential cancer driver mutations^26^. Our results suggest that utilizing flanking region annotations helps boost model performance for separating rare pathogenic variants versus rare (and common) begin variants. In addition, we provide pre-computed MetaRNN scores for all possible human nsSNVs and a Linux executable file for a one-stop annotation of both nsSNVs and nfINDELs.

## MATERIAL AND METHODS

### Training Datasets

ClinVar^27^ dataset files clinvar_20190102.vcf.gz and clinvar_20200609.vcf.gz were downloaded from https://www.ncbi.nlm.nih.gov/clinvar/ under GRCh38/hg38 genome assembly. Variants in the older file were used in the training phase of model development. Next, we prepared separate datasets for point mutations and insertion/deletions (InDels). For SNVs, non-synonymous SNVs (nsSNVs) labeled as ‘Pathogenic’ or ‘Likely pathogenic’ were used as true positives (TPs), and nsSNVs labeled as ‘Benign’ or ‘Likely benign’ were used as true negatives (TNs). Variants with conflict clinical interpretations were removed. Conflict clinical interpretations were defined as one of these scenarios: conflict between Benign/Likely benign and variants of unknown significance (VUS); conflict between Pathogenic/Likely pathogenic and VUS; conflict between Benign/Likely benign and Pathogenic/Likely pathogenic. Rare variants that were absent from at least one of the three datasets (gnomAD^21^, ExAC^20^, and the 1000 Genomes Project^2^) were retained. A further filter removed any nsSNVs that were absent in all three datasets. In the end, 26,517 rare nsSNVs with 9,009 TPs and 17,508 TNs **(Supplementary Table 1**) were used for training. For InDels, the same criteria were applied to obtain TPs and TNs. Additionally, only InDels annotated as non-frameshift (nfINDELs) and have length >1 and < 50 base-pairs were included. A total of 2,057 rare nfINDELs with 1,348 TPs and 709 TNs passed the filtering criteria and were used to train the MetaRNN-indel model **(Supplementary Table 2**).

### Test Datasets

We constructed 8 test sets to evaluate the performance of our SNV based model, namely MetaRNN (Summary in **Supplementary Table 3**) with 24 other methods including MutationTaster^10^, FATHMM^28^, FATHMM-XF^12^, VEST4^9^, MetaSVM^29^, MetaLR^29^, M-CAP^17^, REVEL^4^, MutPred^16^, MVP^8^, PrimateAI^15^, DEOGEN2^14^, BayesDel_addAF^7^, ClinPred^6^, LIST-S2^5^, CADD^3^, Eigen^13^, GERP^30^, phyloP100way_vertebrate^31^, phyloP30way_mammalian, phyloP17way_primate, phastCons100way_vertebrate, phastCons30way_mammalian, and phastCons17way_primate. The first test dataset (rare nsSNV test set, RNTS) was constructed from rare pathogenic nsSNVs (absent from at least one of the three datasets, namely gnomAD, ExAC, and the 1000 Genomes Project, but not absent in all three datasets) added to the ClinVar database after 20190102 and rare nsSNVs absent from at least one of the three datasets but not absent in all three datasets while not reported in ClinVar and matching on genomic location (randomly selected non-pathogenic nsSNVs within 200kb from the pathogenic ones), resulting in 12,406 variants with 6,203 TPs and 6,203 TNs **(Supplementary Table 4**). The second test dataset (rare clinvar-only test set, RCTS) was constructed from recently-curated (after 20190102) ClinVar pathogenic nsSNVs (n = 1,528) and benign nsSNVs (n = 2,389) that were absent from at least one of the three datasets but not absent in all three datasets (**Supplementary Table 5**). The third test dataset (allele-frequency-filtered RNTS, AF-RNTS) was constructed from RNTS with a replacement of the allele count filter with an AF filter that only variants with AF≤0.01 in all three datasets were kept, which resulted in a data set with 5,770 TPs and 5,770 TNs (**Supplementary Table** 6). The fourth test dataset (allele-frequency-filtered RCTS, AF-RCTS) was constructed from RCTS with a replacement of the allele count filter with an AF filter that only variants with AF≤0.001 in all populations were kept, which resulted in 6065 TPs and 3,220 TNs (**Supplementary Table 7**). The fifth test dataset (all-allele-frequency set, AAFS) was constructed from all pathogenic and benign nsSNVs added to the ClinVar database after 20190102, resulting in 29,924 variants with 6,205 TPs and 22,808 TNs **(Supplementary Table 8**). The sixth test set, TP53 test set (TP53TS), was constructed from the TP53 mutation website (https://p53.fr/index.php). Variants with median activity <50 were considered pathogenic, while variants with median activity >=100 are considered benign. After removing variants used in the training set, 824 variants remained with 385 TPs and 439 TNs **(Supplementary Table 9**). The seventh test data set was retrieved from a recent publication (**Supplementary Table 10**) ^26^. The TPs (n = 878) were curated from cancer somatic mutation hotspots and the TNs (n = 1,756) were curated from population sequencing study DiscovEHR ^32^. The Human Gene Mutation Database (HGMD)^24^ is another popular database which provides high-quality disease-associated variants. As the last test set, we retrieved all DM variants (disease mutation; the class of variants in HGMD with highest confidence of being pathogenic) from the HGMD Professional version 2021.01. Only variants reported in dbNSFP as missense were kept. To minimize type I circularity of data, we further removed variants that were reported in the HGMD Professional version 2017. Additionally, variants reported in ClinVar 20200609 as Pathogenic, Likely pathogenic, Benign or Likely Benign were filtered out to explore the generalizability of our score to independently curated disease-causing mutations. These filtered nsSNVs were used as true positives (TP). For true negatives (TN), we used rare nsSNVs that were observed in gnomAD v3 with allele frequency between 0.01 and 0.0001 as a trade-off between their rarity and probability of truly being benign. The number of TP and TN variants were matched which resulted in 45,256 nsSNVs in total (22,628 TP variants and 22,628 TN variants).

For our InDel-based model, namely MetaRNN-indel, the first test dataset was constructed from InDels added to the ClinVar database after 20190102, which resulted in 989 InDels with 491 TPs and 498 TNs **(Supplementary Table 11**). The second test dataset was constructed from HGMD Professional version 2021.01. All the nfINDELs that were not used in training MetaRNN-indel were kept as TP. For TN, rare nfINDELs with AF less than 0.01 were retrieved from gnomAD v2.1.1 as TN, which were then matched on the number of TP. A total of 8,020 nfINDELs (4,010 TP variants and 4,010 TN variants) were collected after filtering.

### Flanking nsSNVs

After obtaining all data sets of target variants, we retrieved their flanking sequences using dbNSFP4.1a^33,34^. Specifically, the variant’s genomic location and the affected amino acid position as to the protein and the affected codon were first identified in dbNSFP. Then, a window size of +/ – 1 codon around the affected codon was identified, and all nsSNVs inside this window were retrieved with maximum length of 9 base-pairs (bps). For a given target mutation, the maximum number of nucleotides on either side is 5 bps (3 bps from one flanking codon and 2 bps from the target codon). To center the input window on target mutation and have a uniform shape for all inputs, we padded the input window to reach an 11 bps window for each target mutation so that there were 5 bps on each side of the target mutation. This window, including the target variants, was used as one input for our model (**Figure 1)**. For each position, multiple alternative alleles may exist. For each annotation at context loci, we calculated the average score across all alleles that would result in a non-synonymous mutation at the locus. This averaged annotation score was then used to represent the locus for that annotation. This setup has the advantage of keeping the most critical context information while limiting the unnecessary noise introduced by having inconsistent order and dimension of alleles at different loci, e.g., some loci may possess three nsSNVs while others may possess only one nsSNV. The target mutation would directly use annotation scores for the observed allele. We assume that these nsSNVs and related annotations can capture the most critical context information with regard to the pathogenicity and functional importance of the amino acids. Please note that we assumed that context nsSNVs provided all the critical information, so we ignored any synonymous mutations in composing the context information. After these steps, the input dimension for the MetaRNN model becomes 11 (bps) by 28 (features, see below).

**Figure 1.**
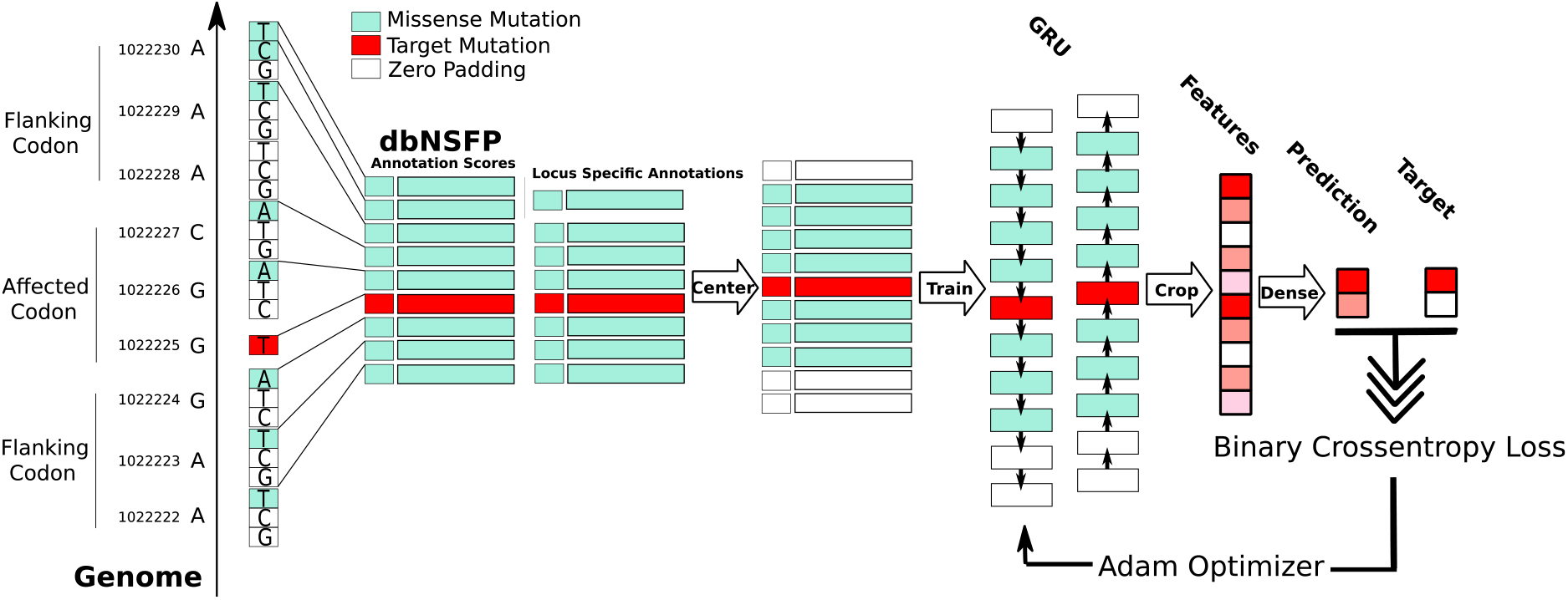
Data preparation and model development steps for MetaRNN.

The same rules to adopting flanking region were applied to InDels with one difference: instead of affecting only one codon, target InDels may directly affect multiple codons simultaneously. Thus, the +/– 1 codon window was defined as the window beyond all the directly affected codons. For deletions, target mutations were those loci deleted by the variant, and their annotations were averaged for each locus. For insertions, since no annotation is available for the inserted nucleotides, we used annotations from loci adjacent to the insertion position as surrogates. Since we focus on short InDels with length <= 48, with 5 bps around each side as context information, the input dimension for the MetaRNN-indel model is 58 (bps) by 28 (features, see below). Again, the synonymous mutations were ignored in composing the context information.

### Feature selection

For each variant, including target nsSNV and flanking region nsSNVs, 28 features were either calculated or retrieved from the dbNSFP database, including 16 functional prediction scores: SIFT^35^, Polyphen2_HDIV^36^, Polyphen2_HVAR, MutationAssessor^11^, PROVEAN^37^, VEST4^9^, M-CAP^17^, REVEL^4^, MutPred^16^, MVP^8^, PrimateAI^15^, DEOGEN2^14^, CADD^3^, fathmm-XF^12^, Eigen^13^ and GenoCanyon^38^; Eight conservation scores including GERP^39^, phyloP100way_vertebrate^31^, phyloP30way_mammalian, phyloP17way_primate, phastCons100way_vertebrate, phastCons30way_mammalian, phastCons17way_primate and SiPhy^40^; Four calculated allele frequency (AF) related scores. The highest AF values across subpopulations of the four data sets from three studies, namely the 1000 Genomes Project (1000GP), ExAC, gnomAD exomes, and gnomAD genomes, were used as the AF scores. All missing scores in the dbNSFP database were first imputed using BPCAfill^41^, and all scores were standardized before feeding to the model for training. Some more recently developed scores were excluded to minimize type I circularity in training our ensemble model, including MPC and ClinPred, which used ClinVar variants in their training process.

### Model development

We applied a recurrent neural network with Gated Recurrent Units^22^(GRU) to extract and learn the sequence information around target variants (**Figure 1**). To select the best-performing model structure, Bayesian Hyperparameter Optimization^42^ was used, and a wide range of model structures was tested. Specifically, the input layer takes an 11×28 matrix as input for MetaRNN and a 58×28 matrix for MetaRNN-indel model. After the bidirectional GRU layer, MetaRNN model cropped out the context information, and only the learned features for the target mutation were kept. This setup can significantly reduce the number of parameters compared to keeping all context features in the subsequent dense layer. Following the same idea, for MetaRNN-indel, the output for the last bidirectional GRU layer only returns the prediction for the final locus (return_sequences parameter was set to false) to limit the number of possible parameters in the following dense layer. The output layer is composed of 1 neuron with a sigmoid activation to model our binary classification problem. A binary cross-entropy loss was used as the loss function, and Adam optimizer^43^ was used to update model parameters through backpropagation^44^. This process used 70% training data for model training and 30% of training data for performance evaluation, so no test datasets were used in this step. Python packages sci-kit-learn^45^ and TensorFlow 2.0 (https://www.tensorflow.org/) were used to develop the models, and KerasTuner (https://keras-team.github.io/keras-tuner/) was adopted to apply Bayesian Hyperparameter Optimization. The search space for all the hyperparameters was shown in **SupplementaryTable 12**. The models with the smallest validation log loss were used as our final models for nsSNV (MetaRNN) and nfINDEL (MetaRNN-indel).

### Compare the Performance of Individual Predictors

As a model diagnosis step, for each model, a permutation feature importance was calculated for each of the individual features. The permutation importance (PI) was calculated as:

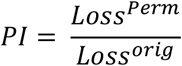

A PI value of 1 means the feature is useless, with higher values indicating more important features. Additionally, SHAP values were calculated to evaluate the direction of impact of each feature on model output ^46^. Using SHAP values, we examined the contribution of each locus in MetaRNN model using the RCTS (n=3,917). Among the 3,917 samples, 90% of them were used as the background set and the rest 10% were used as the set to study. Python library SHAP (https://shap.readthedocs.io/en/latest/index.html) was used to calculate SHAP values and plot visualizations. Using RCTS, we also examined those incorrectly classified instances using SHAP summary plots.

To quantitatively evaluate model performance, we retrieved 39 prediction scores from dbNSFP to compare with our MetaRNN model including MutationTaster^10^, MutationAssessor^11^, FATHMM^28^, FATHMM-MKL^47^, FATHMM-XF^12^, PROVEAN^37^, VEST4^9^, MetaSVM^29^, MetaLR^29^, M-CAP^17^, MPC^18^, REVEL^4^, MutPred^16^, MVP^8^, PrimateAI^15^, DEOGEN2^14^, BayesDel (AF and noAF models)^7^, ClinPred^6^, LIST-S2^5^, LRT^48^, CADD (raw and hg19 models)^3^, DANN^49^, Eigen (raw and PC models)^13^, GERP^30^, Polyphen2 (HVAR and HDIV)^50^, SIFT4G^51^, SiPhy^40^, GenoCanyon^52^, fitCons (integrated)^53^, phyloP (100way_vertebrate, 30way_mammalian and 17way_primate)^31^ and phastCons (100way_vertebrate, 30way_mammalian and 17way_primate)^54^. The corresponding rankscores was retrieved for each of these 39 annotation scores to facilitate comparison. For the MetaRNN-indel model, four popular methods were compared, including DDIG-in^55^, CADD^3^, PROVEAN^37^, and VEST4^56^. For the ClinVar holdout test set, all these four methods were compared with MetaRNN-Indel. For the HGMD test set, VEST4 was removed from comparison since it used HGMD InDels during training, and we do not have access to an older version of HGMD with InDels to exclude these variants. For both test data sets, LiftOver was used to convert hg38 genomic coordinates to GRCh37/hg19 for DDIG-in and PROVEAN. DDIG-in scores were retrieved from https://sparks-lab.org/server/ddig/. VEST4 indel scores were retrieved from http://cravat.us/CRAVAT/. The CRAVAT format was used, and each indel mutation was assumed to be located on both + and – strands. For PROVEAN indel, the scores were retrieved from http://provean.jcvi.org/genome_submit_2.php?species=human. The CADD scores v1.6 under GRCh38 assembly were obtained from http://provean.jcvi.org/genome_submit_2.php?species=human. We plotted receiver operating characteristic (ROC) curves and calculated the area under the ROC curve (AUC) for each method being compared using our test datasets. Python package matplotlib (https://matplotlib.org/) was used to plot ROC curves, and Python package sci-kit-learn was used to calculate AUC scores.

### Development of MetaRNN and MetaRNN-indel Linux executables

To facilitate custom annotations with user-provided VCF files, we created a Linux executable using Python library PyInstaller (https://www.pyinstaller.org). This library freezes Python applications into stand-alone executables, effectively solving issues related to incompatible environments and missing dependencies. Briefly, the executable includes the following steps to make final predictions of MetaRNN and MetaRNN-indel scores. First, it takes as input a VCF file that includes candidate variants. Second, ANNOVAR^57^ was used to annotate these variants, and only nsSNVs and nfINDELs were kept. Third, for nsSNVs, the executable will extract MetaRNN predictions from our database of all pre-calculated nsSNVs; for nfINDELs, target mutation and its context mutations will be first identified, and all required annotations will be retrieved from dbNSFP^33^ and the MetaRNN-indel model will be used to make predictions on these user-provided nfINDELs. Lastly, two output files will be generated for nsSNVs and nfINDELs separately.

## Results

### Allele Frequencies as Crucial Features in Separating Pathogenic Mutations

MetaRNN and MetaRNN-indel models took an ensemble method that combined 24 individual prediction scores and four allele frequency (AF) features from the 1000 Genomes Project (1000GP)^58^, ExAC^20^, and gnomAD^21^. As shown in **Figure 2A**, most of the component conservation scores and ensemble scores showed moderate to strong correlations (correlation coefficient between 0.4 and 1). However, MutationTaster^10^ and GenoCanyon^38^ showed a weak correlation with all other features. Since most SNVs are not observed in multiple populations (AF=0), correlations between different AF features are also strong (>0.8). AF features showed a weak correlation with all other individual predictors implying that previous annotation scores have not fully exploited such allele frequency information. This observation is also supported by the permutation feature importance analysis (**Figure 2B**). The most important features are two exome AF features, followed by two whole-genome AF features. Some functional predictors showed higher importance compared with others, such as VEST4^9^, MutationTaster^10^, and MutPred^16^. This observation is in concordance with the previous observations^6^ highlighting the importance of population AF data in inferring the functional importance of nsSNVs.

**Figure 2.**
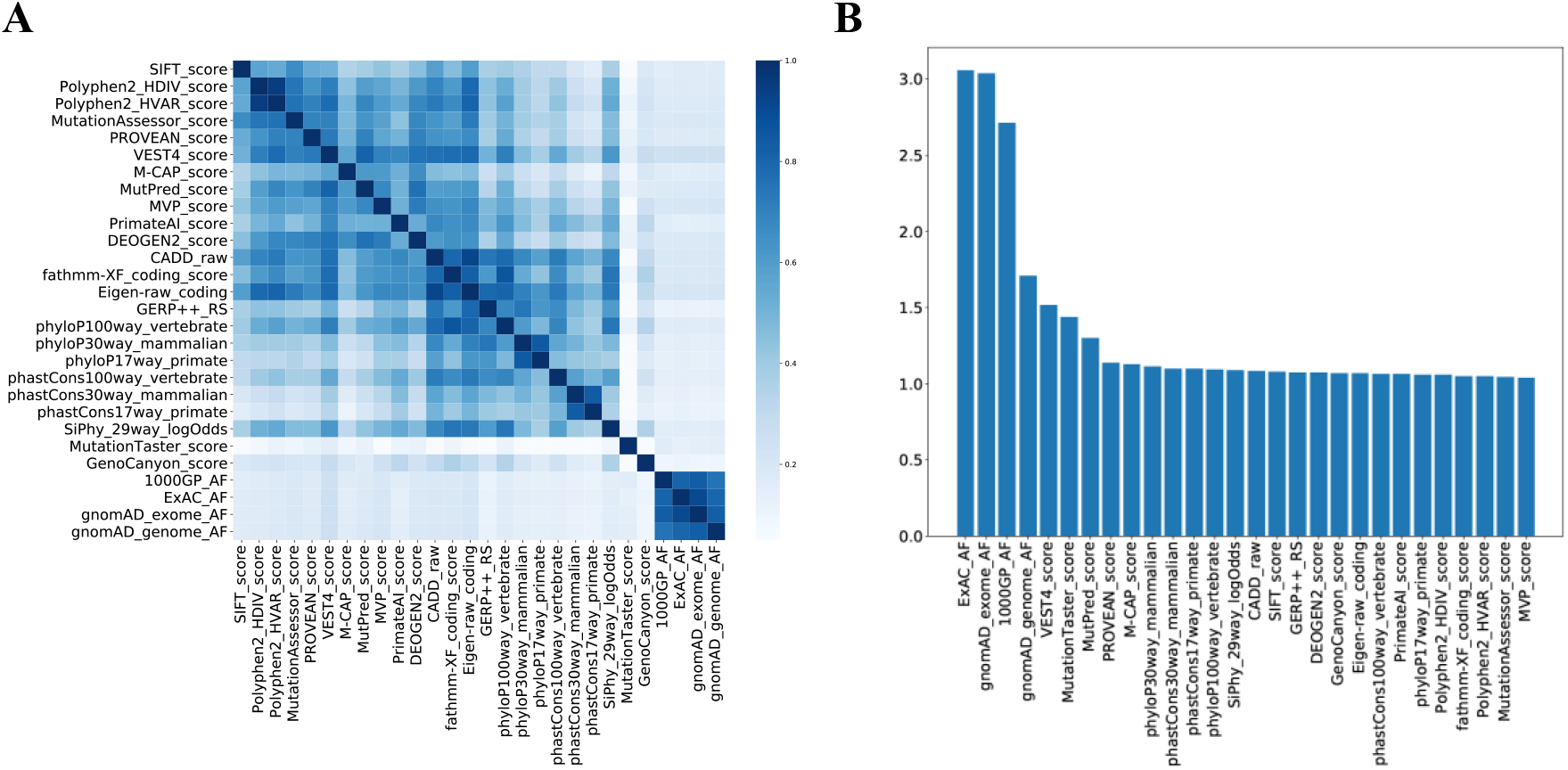
Features used to train MetaRNN and MetaRNN-indel. A. Correlation between features used to train MetaRNN. B. Feature importance for all features used in the MetaRNN model.

### Performance Comparison of MetaRNN to Other Predictive Algorithms Using ClinVar

As the major goal of MetaRNN model is to separate between rare pathogenic from rare benign nsSNVs, we constructed two rare nsSNV test sets (RNTS and AF-RNTS, see **METHODS**) that were composed of rare pathogenic ClinVar nsSNVs after release 20190102 and location-matched rare benign nsSNVs from gnomAD, ExAC and the 1000GP. The RNTS (*n* = 12,406) test set was filted based on the presence/absence of variants across different populations whereas the AF-RNTS (*n* = 11,540) was filtered based on more fine-grained allele frequency data. For the RNTS set, MetaRNN achieved the best performance with an area under the ROC curve (AUC) equals 0.9100 in separating these rare nsSNVs, followed by BayesDel_addAF^7^ and REVEL^4^ (selected comparisons with eight tools are available in **Figure 3A**; all comparisons with 24 tools are available in **S. Figure 9**). It has been reported that computational tools tend to overestimate the number of pathogenic mutations (i.e., high sensitivity and low specificity)^59,60^. Consequently, we then examined the models’ specificity at 95% sensitivity. The MetaRNN model achieved the best specificity (0.6105) as to correctly identifying rare benign nsSNVs, followed by REVEL (0.5037) and BayesDel_noAF (0.4936). For the AF-RNTS set, MetaRNN outperformed all competitors with an AUC of 0.9322, followed by BayesDel (AUC=0.9235) and ClinPred^6^ (AUC=0.9205) (**S. Figure 2**). The MetaRNN model again achieved the best specificity (0.6988) at 95% sensitivity, followed by ClinPred (0.647) and BayesDel_addAF (0.6415). Similarly, MetaRNN showed the best performance using the RCTS (AUC=0.9156) and AF-RCTS (AUC=0.9446) test sets (**S. Figure 8** and **S. Figure 9**).

**Figure 3.**
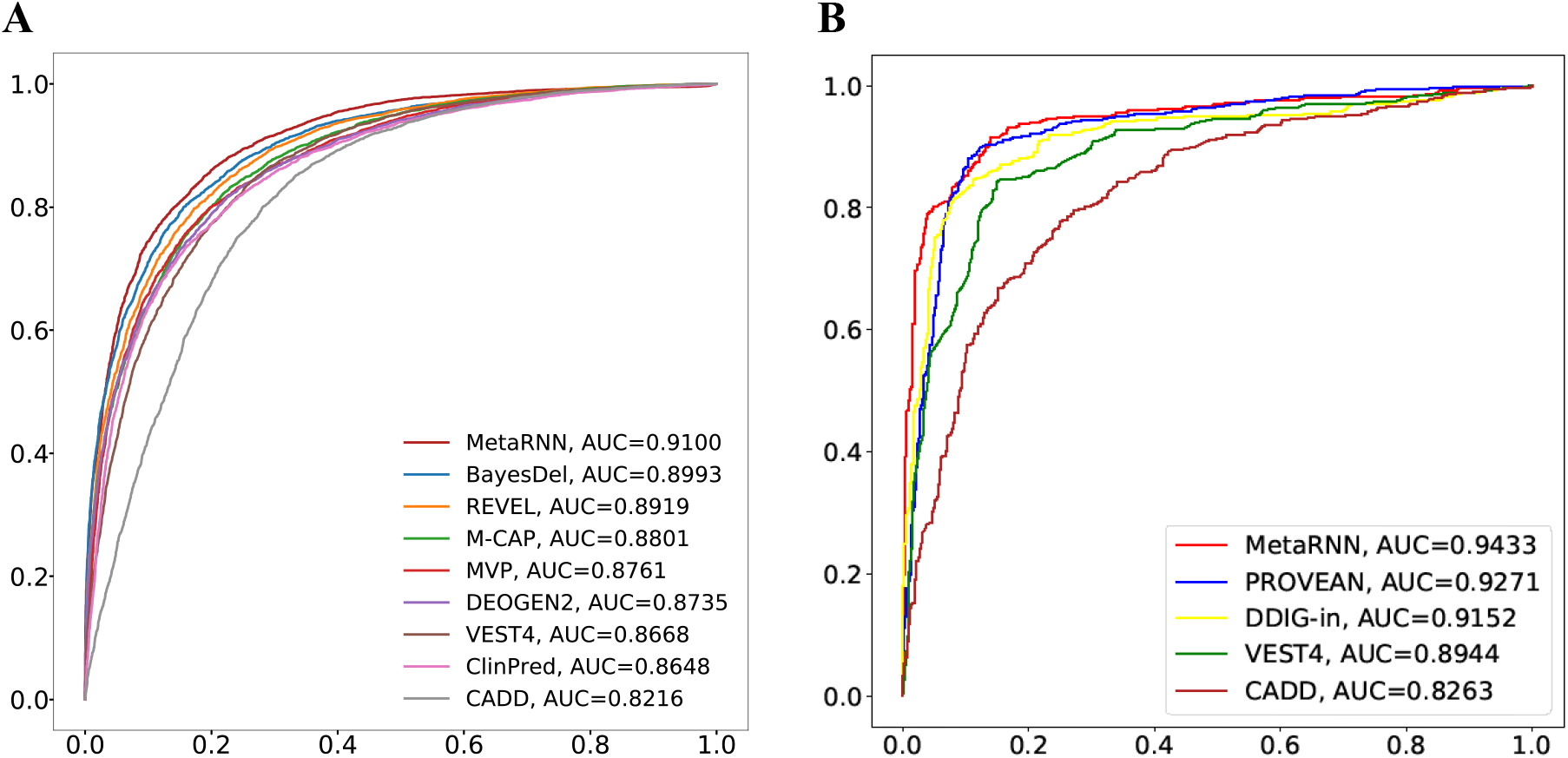
Comparisons of MetaRNN and MetaRNN-indel with other prediction tools. **A**: Performance comparison of MetaRNN and 8 other nsSNV prediction tools using the rare nsSNV test set (RNTS). **B**: Performance comparison of MetaRNN-indel and other nfINDEL prediction tools using ClinVar nfINDELs.

To evaluate the performance of MetaRNN in separating all ClinVar reported nsSNVs, we constructed an all-allele-frequency set (AAFS) comprised of all available ClinVar pathogenic SNVs and benign SNVs (rare+common) that are not used for model development. As shown in **S. Figure 4**, using AAFS as benchmarking dataset, MetaRNN outperforms all competitors with an AUC of 0.9862. The second-best model is ClinPred (AUC=0.9841) and followed by BayesDel (AUC=0.9759). In general, ensemble methods and functional predictors outperform conservation-based methods in this comparison. In addition, MetaRNN showed improved performance under both benchmark settings with and without an AF filter for the inclusion nsSNVs.

### Investigating Generalizability of MetaRNN to Different Disease and Functional Databases

To explore the generalizability of our model on disease-causing nsSNVs curated with different standards, we retrieved disease-causing mutations (DM) from the HGMD Professional v.2021.01 ^24^ as true positives (n = 22,628) and matched rare nsSNVs observed in gnomAD v3 with allele frequencies between 0.01 and 0.0001 as true negatives (n = 22,628). Only variants reported in dbNSFP^33,34^ as missense were kept. To minimize type I circularity of data, we further removed variants that were reported from an older version HGMD Professional database (v. 2017). Additionally, variants reported in ClinVar 20200609 as “Pathogenic”, “Likely pathogenic”, “Benign” or “Likely Benign” were filtered out. Using this test dataset, MetaRNN still outperformed other competitors with an AUC equals 0.9689 (**S. Figure 7**). TP53 is one of the most well-studied human genes, with its functional impact linked to tumor suppression^61^. Using results from a TP53 mutagenesis assay (*n* = 824), we showed that MetaRNN provides the best estimations for results from such functional experiments with an AUC equals 0.8074 (**S. Figure 5**). Additionally, we collected a dataset of cancer somatic mutations from a recent study and showed that both MetaRNN and BayesDel showed the best performance in separating potential driver mutations from potentially benign variants observed in populations (**S. Figure 6**) ^26^. These results highlighted MetaRNN’s increased ability relative to those of the other methods to separate not only rare pathogenic mutations from rare benign ones but also mutations with various degrees of functional importance across different disease pathways.

### MetaRNN-indel Outperformed Competitors in Identifying Pathogenic nfINDELs

To examine the performance of MetaRNN-indel, we first curated a test dataset that was composed of rare pathogenic ClinVar nfINDELs after release 20190102 (*n* = 989). MetaRNN-indel outperformed all competitors in ranking nfINDELs with an AUC equals to 0.9433 (**Figure 3B**), including two methods, VEST^56^ and CADD^3^, that showed good performance for nsSNVs. The second test dataset was constructed from HGMD Professional version 2021.01. All the nfINDELs that were not in the training set of MetaRNN-indel were used as the pathogenic set. For benign set, rare nfINDELs with AF less than 0.01 were retrieved from gnomAD v2.1.1, then matched the number of pathogenic variants. A total of 8,020 nfINDELs (4,010 pathogenic variants and 4,010 benign variants) were collected after filtering. MetaRNN-indel still outperformed other scores with an AUC of 0.8491, followed by PROVEAN (AUC=0.7951) (**S. Figure 7**).

### MetaRNN Showed Improved Interpretability of Variant of Unknown Significance

To explore the interpretability and usability of the proposed models, we first predicted scores for all nsSNVs in ClinVar that showed conflicting clinical interpretations (*n* = 20,337). These nsSNVs represent an essential class of variants with unknown significance (VUS) according to the ACMG-AMP guidelines^62^. The ability to distinguish and interpret VUSs is crucial to the clinical application of the proposed score. A score that shows sufficient dispersion enables further identification of relevant candidate variants. Additionally, these conflicting VUS variants are of interest with some evidence of being either pathogenic or benign. Among these variants, 15,788 (77.6%) showed conflicting interpretations between benign/likely benign and unknown significance (“benign conflicting group”), whereas 4,110 (20.2%) showed conflicting interpretations between pathogenic/likely pathogenic and unknown significance (“conflicting pathogenic group”). Based on the fact that the benign conflicting group had approximately four times more variants than the conflicting pathogenic group, we expect that variant prediction tools should reflect this observation. While other scores either showed little change in the distribution across their predictions (e.g., CADD^3^, VEST^9^, REVEL^4^) or potentially underestimated the proportion of VUSs at the extremes (BayesDel^7^), MetaRNN’s predictions showed a score distribution that fit these assumptions (**Figure 4A**), which peaked at the extremes of its score range and had approximately four times more extreme benign predictions than extreme pathogenic predictions.

**Figure 4.**
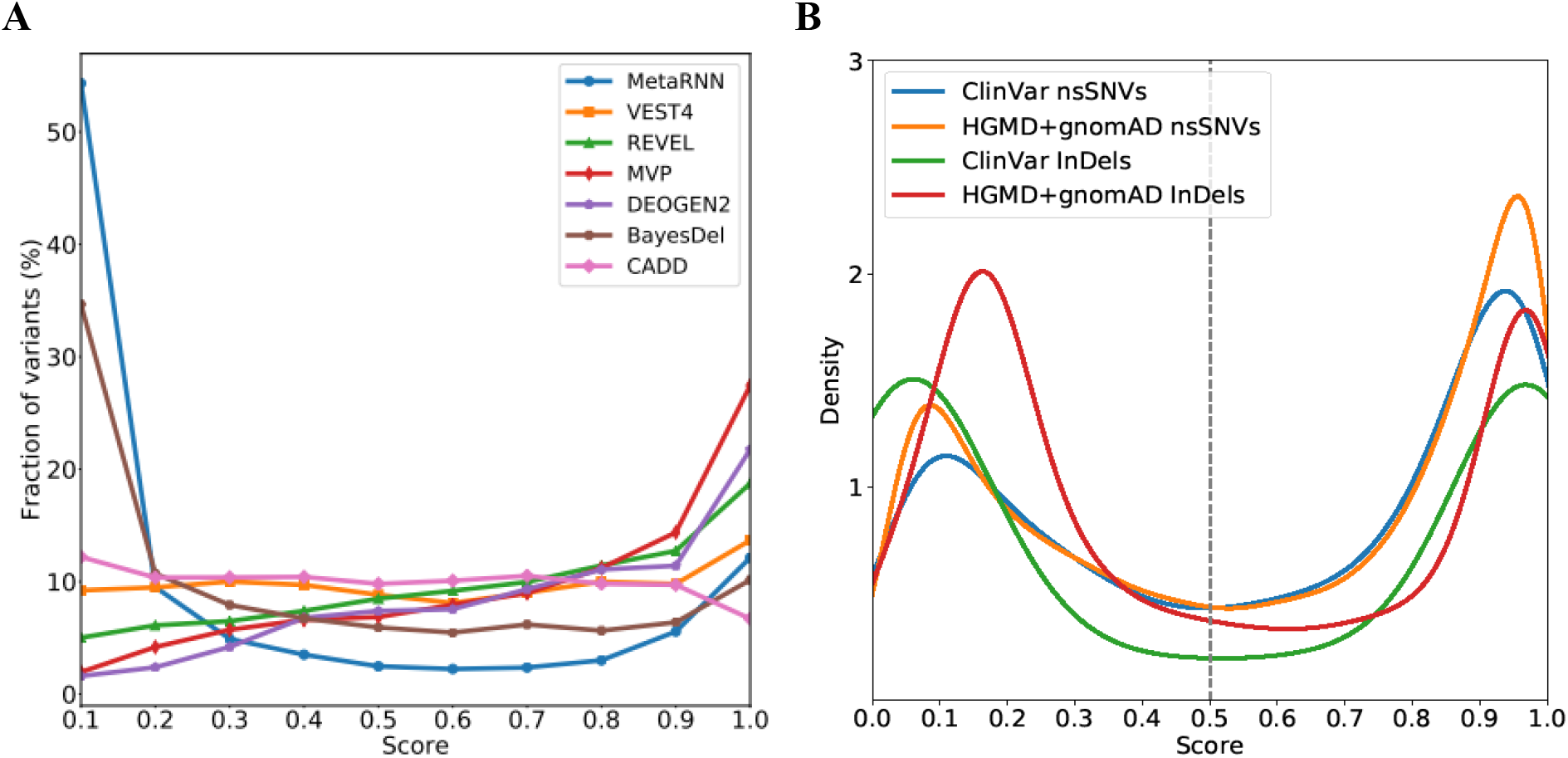
MetaRNN and MetaRNN-indel score distributions in test data sets. **A**: Score distribution for ClinVar variants of unknown significance (VUSs). **B**: Score distributions of MetaRNN and MetaRNN-indel predictions on matched test datasets where the number of pathogenic and benign mutations are the same.

### Compatibility of MetaRNN and MetaRNN-Indel Scores

Additionally, we explored the score distributions for nsSNVs and nfINDELs used in testing the respective MetaRNN models. A clear bimodal distribution was observed for both MetaRNN and MetaRNN-indel predictions (**Figure 4B**). Based on a cutoff value of 0.5 as inherited by the sigmoid activation function used in the MetaRNN models, pathogenic nsSNVs and nfINDELs can be effectively separated from benign ones. These observations have two implications. First, using a cutoff of 0.5 is in accordance with the interpretation of the scores as probabilities where a score greater than 0.5 can be categorized as a higher probability of being pathogenic, and a score less than 0.5 can be categorized as a higher probability of being benign. Second, with a shared cutoff value and similar distributions for nsSNV and nfINDEL scores across independent test datasets, we show predictions from our two models, namely MetaRNN and MetaRNN-indel, are comparable. This feature can effectively help increase the power of genotype-phenotype association studies and related gene-set association analyses. It can also help fine-mapping to pinpoint the exact causal mutations in coding sequences.

### Context Information as Important Contributors to Model Performance

Lastly, to show that our MetaRNN model, which adopted flanking sequence information, can provide additional predictive power, we first trained another feedforward neural network model using only dense layers with feedforward connections (MetaRNN.feedforward). The same KerasTuner hyperparameters were applied, such as the maximum number of trials and the number of hidden layers. Additionally, we checked if limiting training data to only those rare nsSNVs (MAF≤0.01 in all populations; n=18,070) could improve the model’s performance in separating rare pathogenic from rare benign nsSNVs (MetaRNN.rare). The comparison showed that our original MetaRNN model outperforms all these alternative model structures (**S. Figure 10**). Models which adopted flanking information, MetaRNN (AUC=0.9322) and MetaRNN.rare (AUC=0.9277), outperformed the model structure that used only target site information (MetaRNN.feedforward, AUC=0.9116)—additionally, limiting training data to rare variants negatively impacted model performance with a slightly decreased AUC value.

To further explore locus-specific contributions of our model, we calculated SHAP (SHapley Additive exPlanations) values ^46^. Unlike conventional permutation-based feature importance, SHAP can measure the direction of impact on an outcome. Therefore, it provides better interpretability of the model. As shown in **Figure 5A**, while the target mutation showed the highest importance, as represented by the magnitude of SHAP values along the x-axis, surrounding loci except for positions 1, 2, and 11 showed a noticeable contribution to the model prediction. Using SHAP values, we also investigated misclassified nsSNVs to identify the source that drives the incorrect classification. Fifty-five false-positive predictions were studied where benign variants were wrongfully classified as pathogenic (**Figure 5B**). The grey vertical line indicates the SHAP value with no impact on the model prediction (SHAP value equals 0), while points to the right of the line indicate that the feature scores can positively impact model prediction, i.e. bias the prediction to be close to 1 (pathogenic). From the figure we can learn that the feature scors at the target position (position 6) drive the prediction to be pathogenic, even when the feature scores have low values as represented by the blue points. On the other hand, features at surrounding positions, such as position 7 and position 9, can still provide balanced information that when individual predictors have low values (blue points), they can still correctly contribute to an overall negative prediction (benign). A similar trend was observed for cases when the model gave false-negative predictions (**Figure 5C**). These observations suggest that features from surrounding loci could provide useful information for pathogenicity prediction as well as regularization to compensate for the potential biased prediction contributed by the target locus.

**Figure 5.**
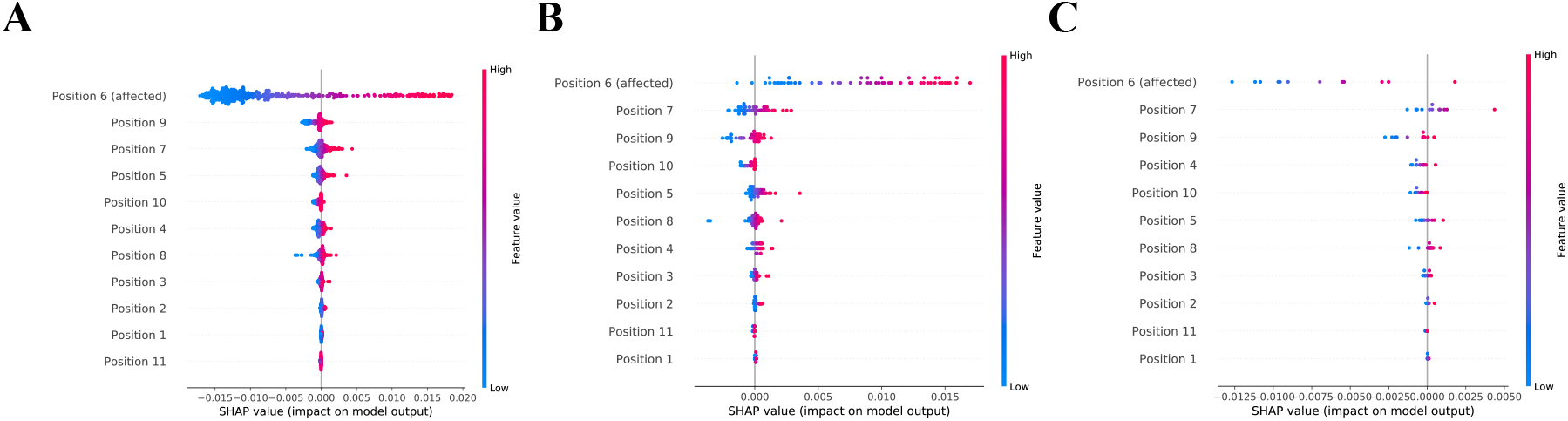
SHAP values for MetaRNN model using RCTS dataset. **A**: SHAP values for 10% of all predictions in RCTS (n=392). **B**: SHAP values for false positve predictions in RCTS (n=55). **C**: SHAP values for false negative predictions in RCTS (n=12).

## Discussion

This study proposed two supervised deep learning models to distinguish pathogenic nsSNVs and nfINDELs from benign ones effectively. Compared to other competitors, MetaRNN showed improved overall AUC and specificity across various test data sets. The improved performance can be attributable to several factors. First, allele frequencies were used as features. Some evidence and observations from our study have shown that population allele frequency can provide valuable information to help to separate pathogenic from benign mutations ^6,7,12^. Our training data, which removed only those “easy” benign nsSNVs observed in all populations, seem to be a good trade-off between posing a difficult enough training set for the model to learn useful information from and preserving valuable information from AF features. Second, information from nsSNVs franking the target mutation helps predict the pathogenicity of the target mutation. Our results indicate that incorporating annotations from context nsSNVs, which are previously neglected by other computational tools, can improve model performance by providing additional context information and providing regularization to minimize overfitting. In future development, different modes of inheritance of diseases and penetrance can be incorporated into model development, making AF information even more useful.

Improved score interpretability is another highlight of the models. As clinical laboratories report candidate mutations mainly based on the ACMG-AMP guidelines^62^, reliable and powerful computational approaches can be a cost-effective way of providing supporting evidence to variant interpretation (such as PM4 and PM5 criteria from the guidelines). By correctly assigning more VUSs into functional groups (pathogenic/benign), more de novo mutations or mutations with insufficient evidence are likely to be interpreted, leading to an improved diagnostic rate in rare Mendelian disorders.

As illustrated previously, MetaRNN and MetaRNN-indel scores are compatible, which filled another gap by providing a one-stop annotation score for both types of mutations. This improvement is expected to be applicable across various settings, such as integrated (nsSNV+nfINDELs) rare-variant burden test for genotype-phenotype association. Even though NGS-based studies such as whole-exome sequencing studies are designed to detect rare genetic mutations, their ability to systematically assess rare genetic variants’ contribution to human diseases and phenotypes still lags behind due to insufficient power. This is contributable to both the low AF of the detected mutations and relatively low sample sizes compared to genotype-based studies^63^. Using computational prediction scores as weights in burden tests is able to increase the power of such studies^64^. The power increase shall be more prominent when nsSNVs and nfINDELs are analyzed in an integrated fashion instead of being analyzed separately.

We provide predictions for all potential nsSNVs (∼86 million) in the dbNSFP^33,34^ database for rapid and user-friendly analysis and a stand-alone Linux executable for the Linux environment for nfINDEL (and nsSNV) predictions. The executable takes a standard VCF file as input and provides variant pathogenicity scores in a transcript-specific manner as output (supported by ANNOVAR^57^). The average prediction time for a single insertion/deletion is approximately 0.2 seconds, which can support timely large-scale predictions.

In conclusion, with improved prediction accuracy, score interpretability, and usability, MetaRNN and MetaRNN-indel will provide more accessible and accurate interpretation of rare VUSs for exome-sequencing-based Mendelian disease studies and integrated (nsSNV+nfINDELs) burden tests for common disease studies.

## Supplementary Data

Supplementary data include 1) 12 supplementary tables including training and testing data used in the study; 2) additional results of 10 figures.

## Acknowledgements

The research was supported by the National Human Genome Research Institute grant 1R03HG011075 to X.L.

## Web Resources

dbNSFP v4.1, https://sites.google.com/site/jpopgen/dbNSFP

ANNOVAR, https://annovar.openbioinformatics.org/en/latest/

KerasTuner, https://www.tensorflow.org/tutorials/keras/keras_tuner

Tensorflow, https://www.tensorflow.org/

PyInstaller, https://www.pyinstaller.org/downloads.html

ClinVar, https://www.ncbi.nlm.nih.gov/clinvar/

HGMD, http://www.hgmd.cf.ac.uk/

## Data and Code Availability

All pre-computed nsSNVs scores as well as the Linux executable file is available at http://www.liulab.science/MetaRNN.

