## Supplementary Figures for "MetaRNN: Differentiating Rare Pathogenic and Rare Benign Missense SNVs and InDels Using Deep Learning"

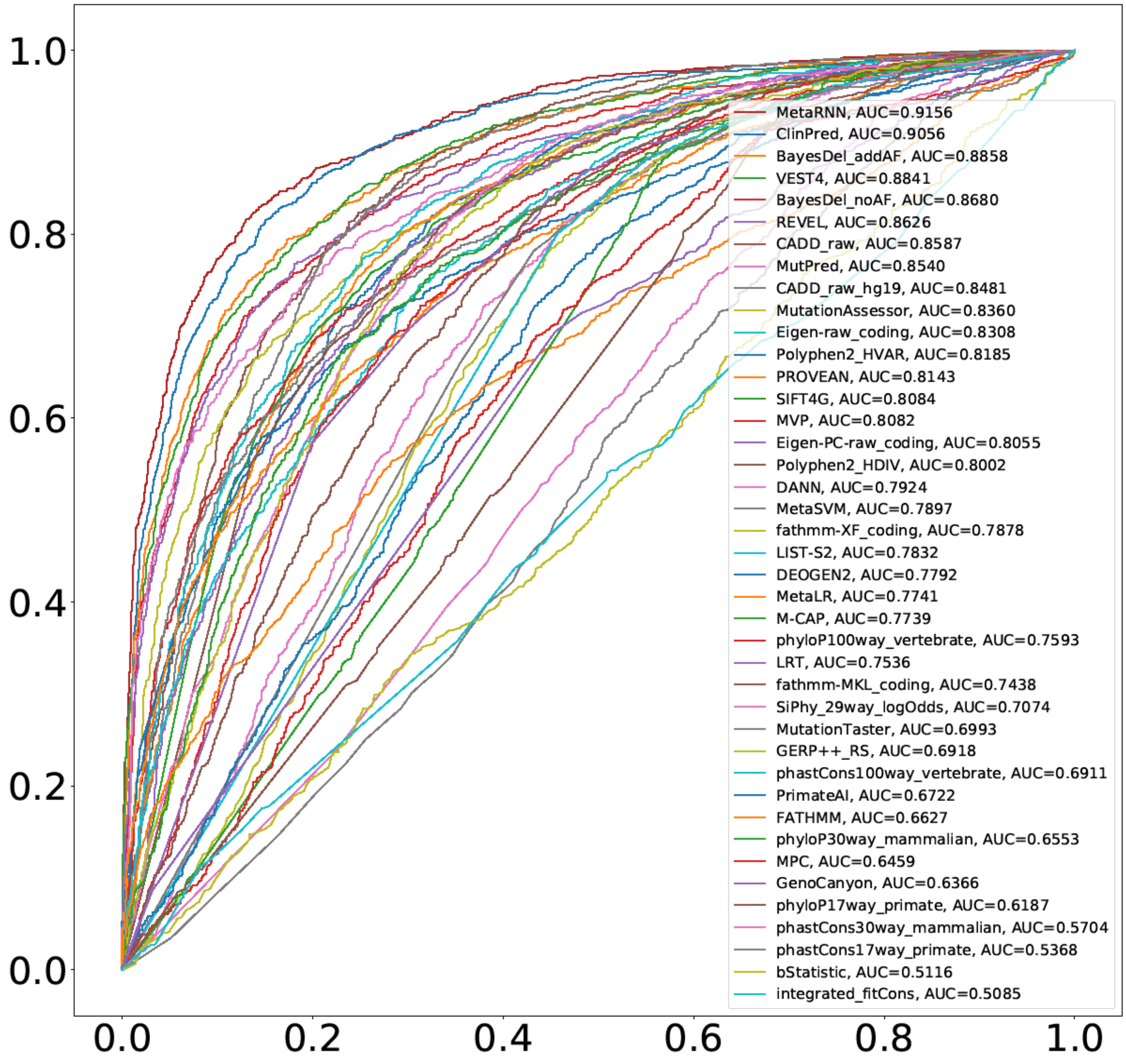


**S. Figure 1. Performance of different methods benchmarked using rare ClinVar-only test set (RCTS).**

### Supplementary Note

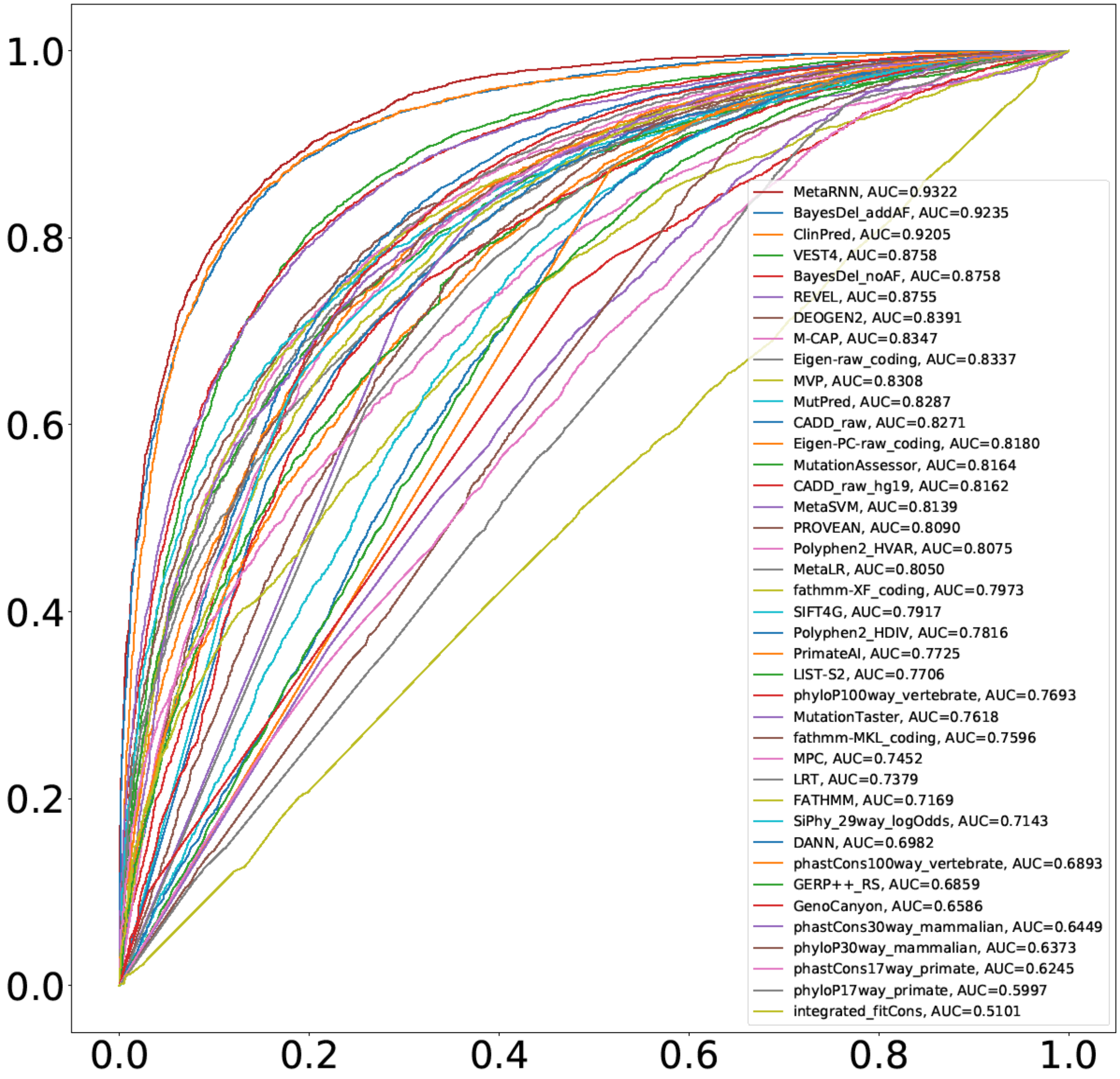


**S. Figure 2. Performance of different methods benchmarked using the allele-frequency-filtered rare nsSNV test set (AF-RNTS).**

### Supplementary Note

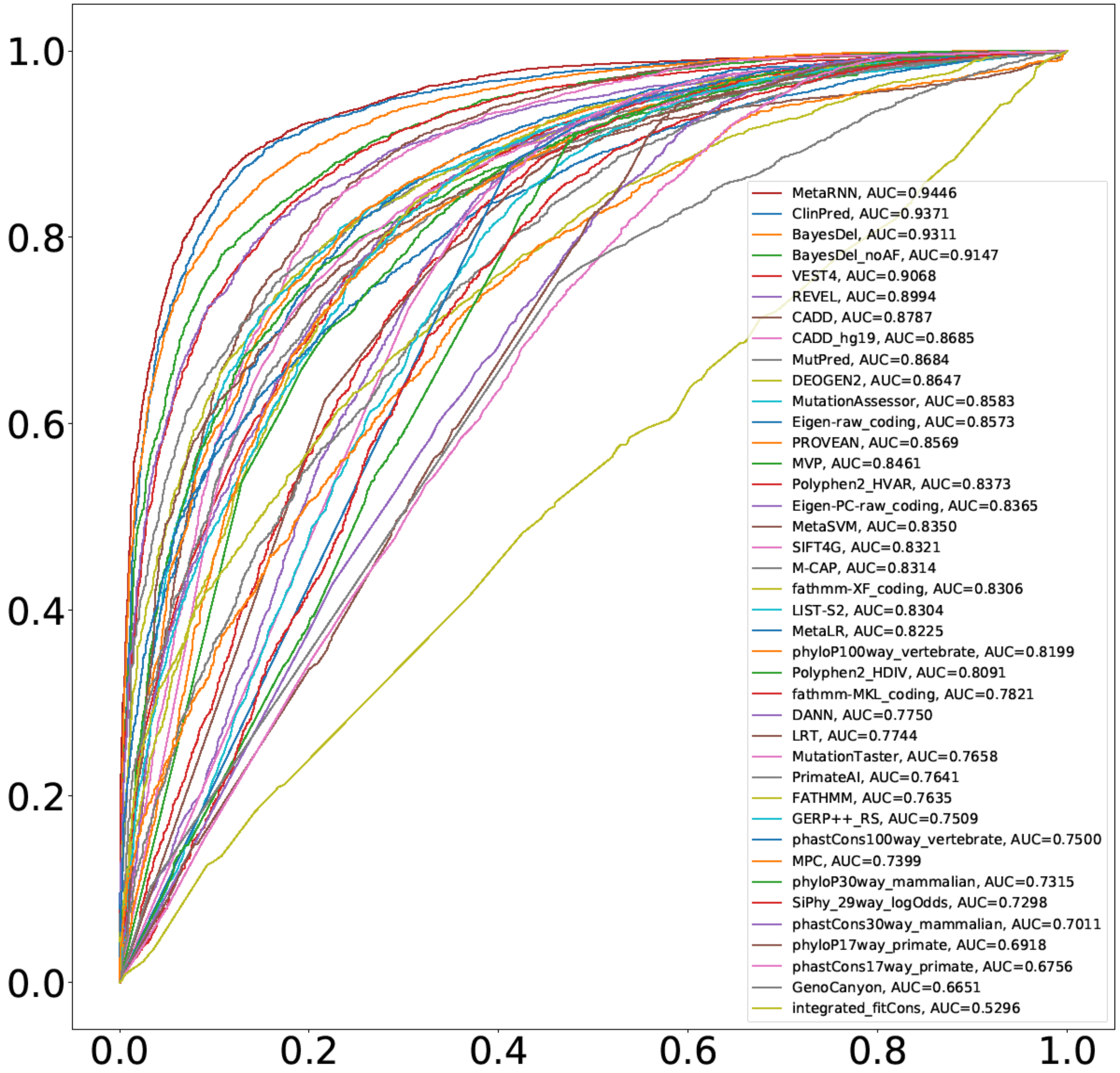


**S. Figure 3. Performance of different methods benchmarked using allele-frequency-filtered rare ClinVar-only test set (AF-RCTS).**

### Supplementary Note

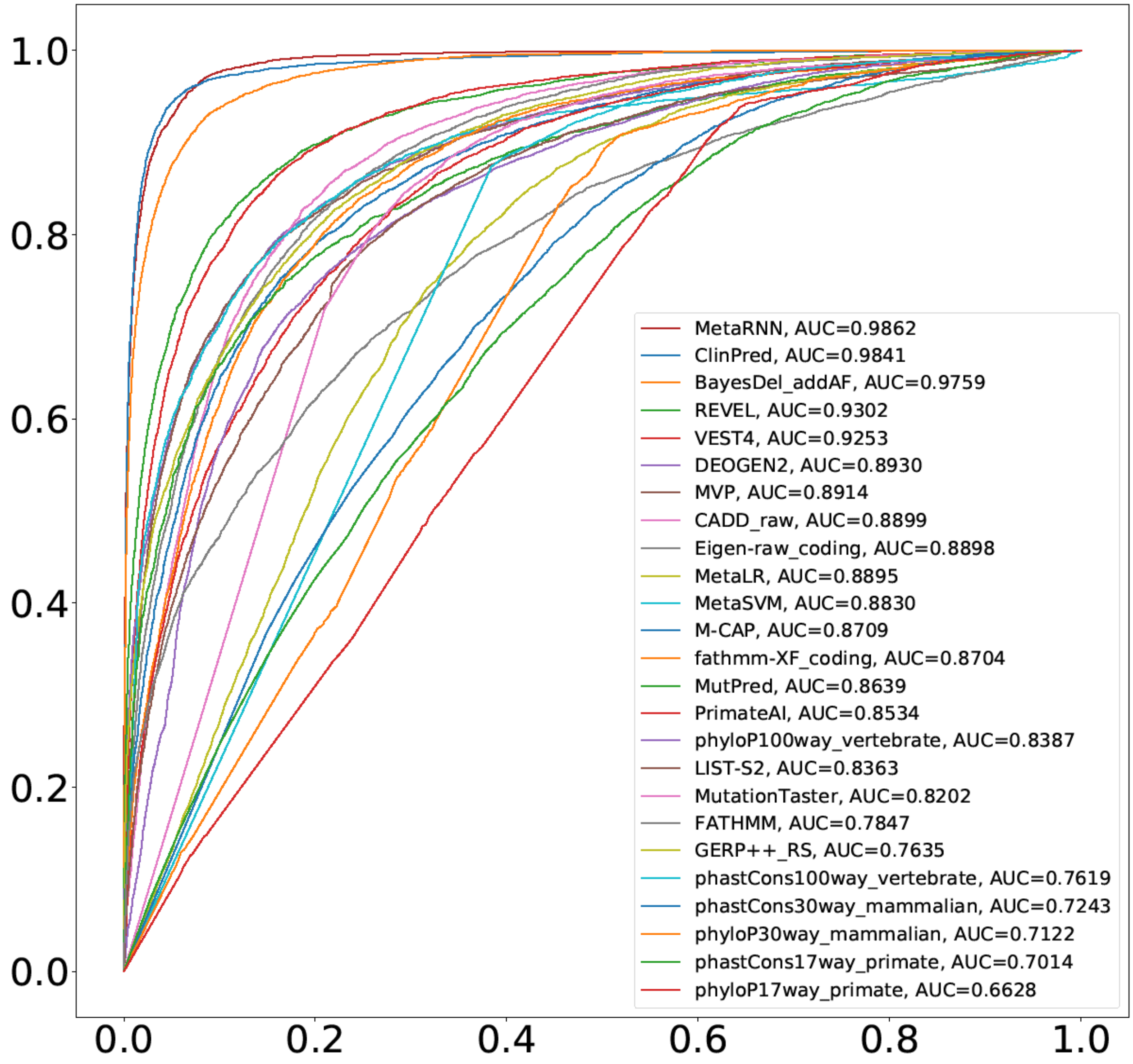


**S. Figure 4. Performance of different methods benchmarked using the all-allele-frequency set (AAFS).**

### Supplementary Note

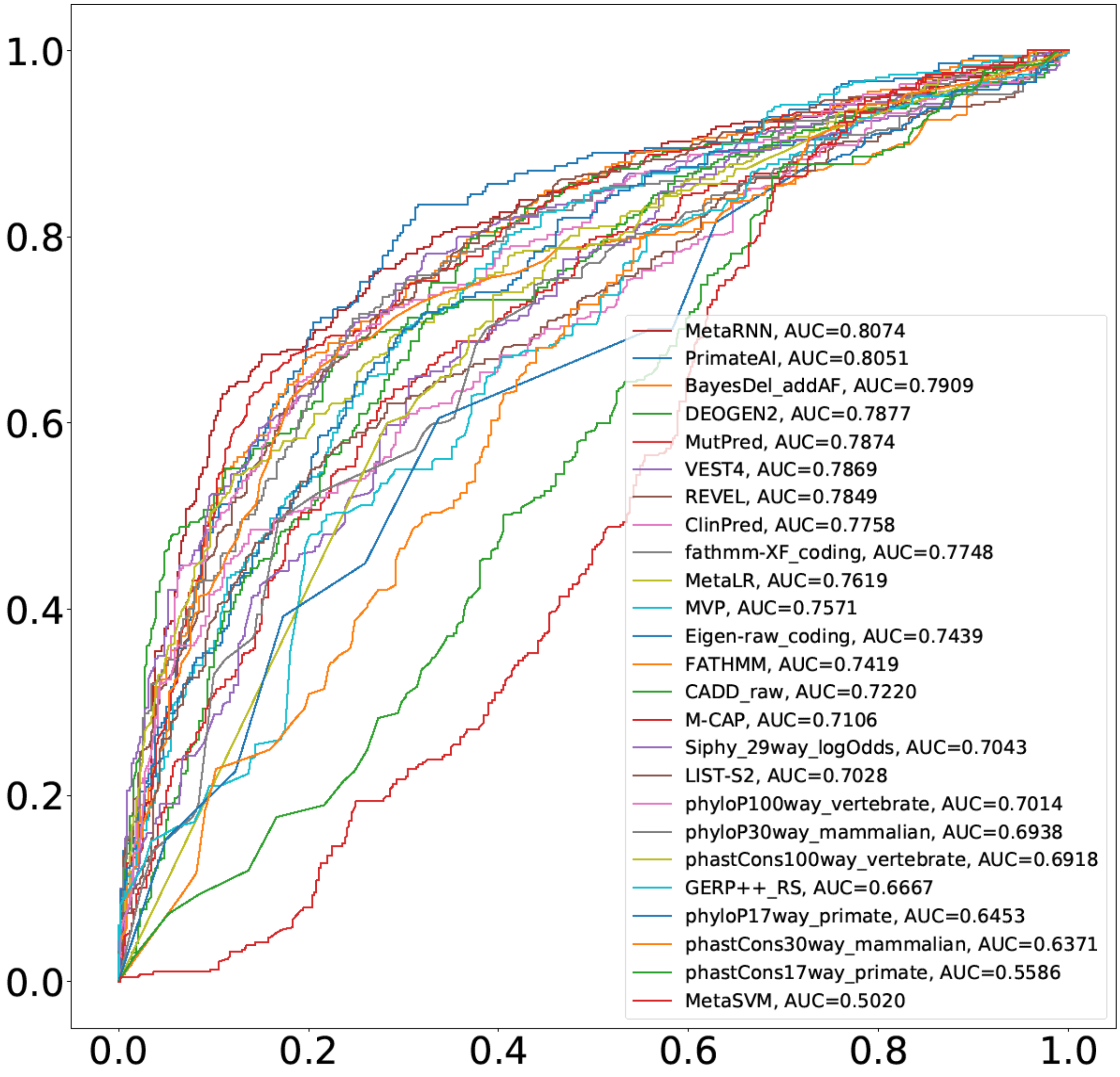


**S. Figure 5. Performance of different methods benchmarked using TP53 test set (TP53TS).**

### Supplementary Note

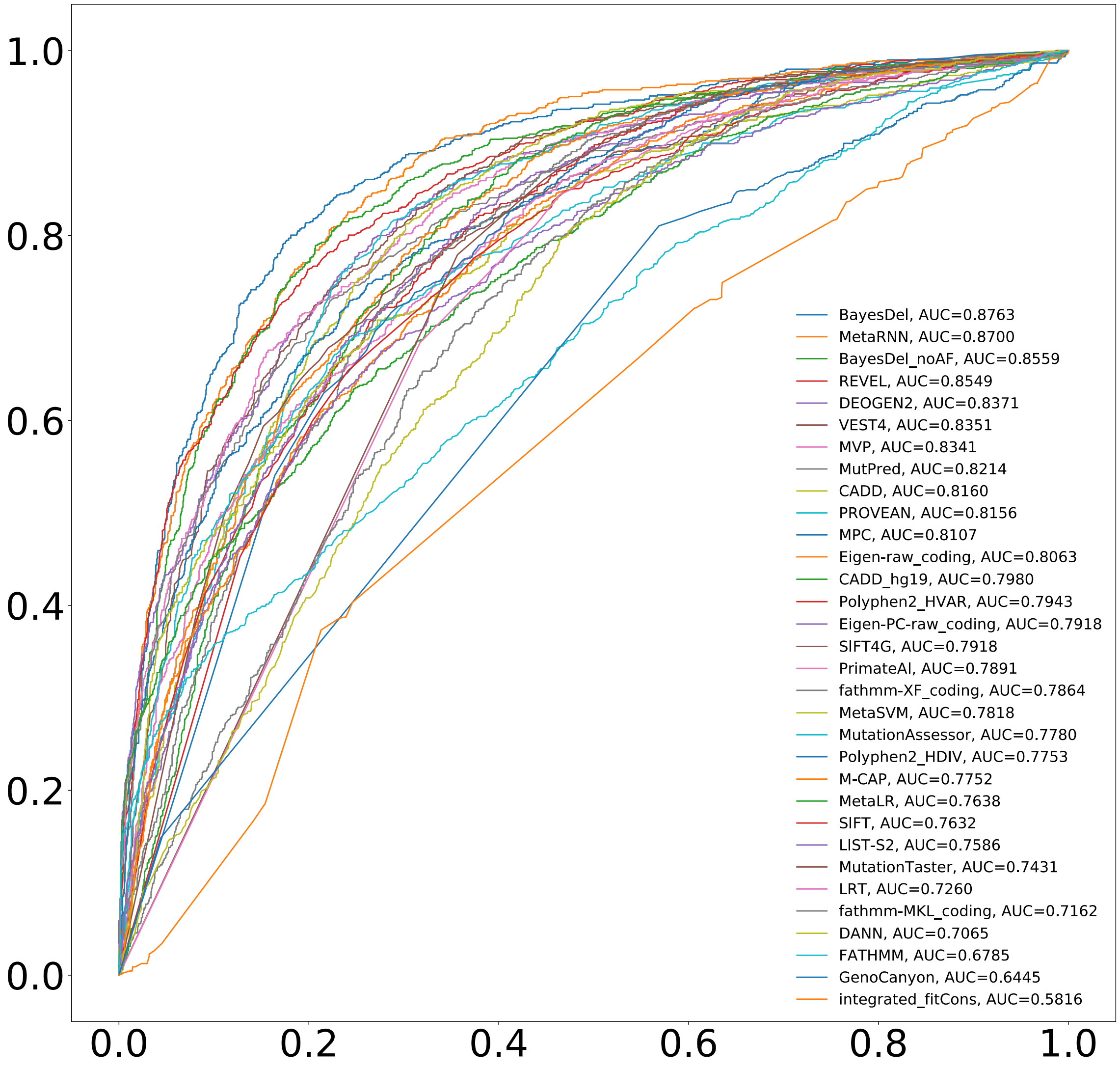


**S.Figure 6. Performance of different methods benchmarked using cancer somatic hotspot mutations as TPs and population sequencing mutations from DiscovEHR as TNs.**

### Supplementary Note

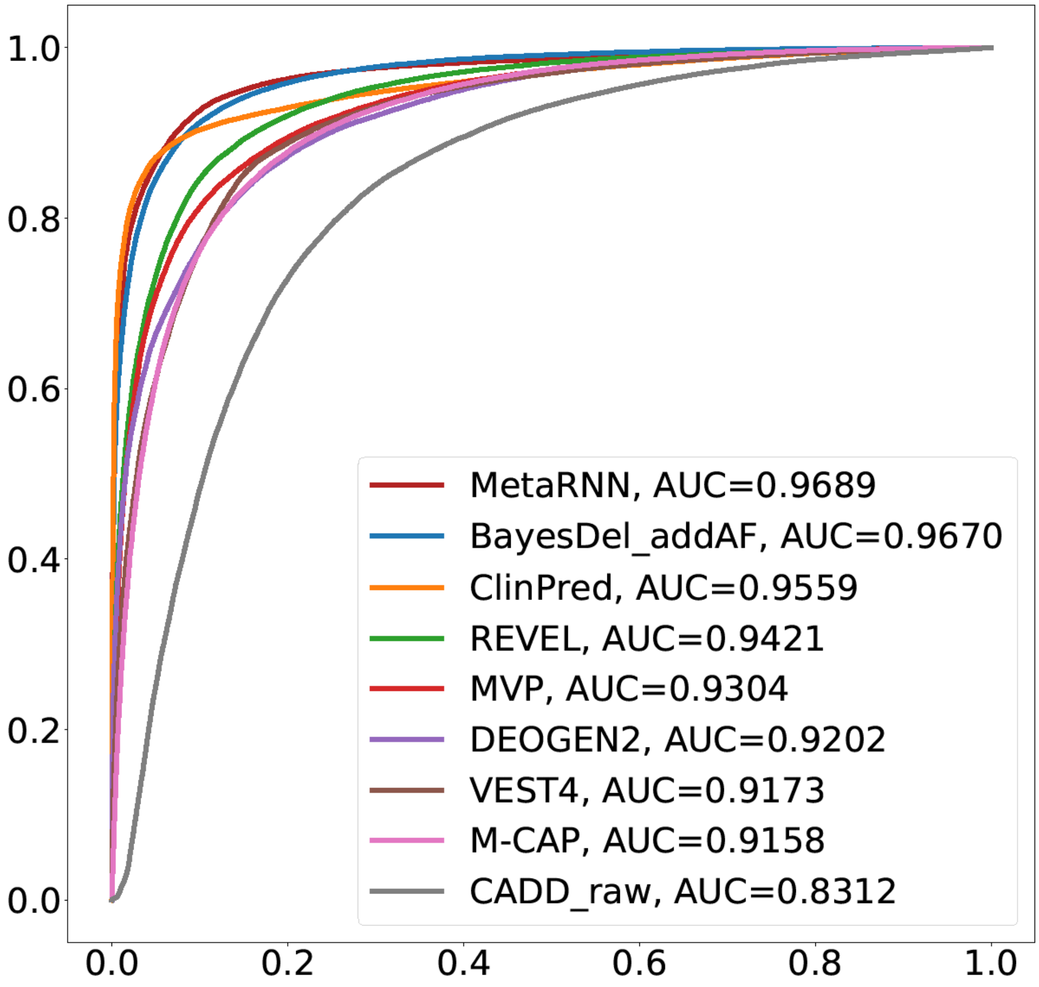


**S. Figure 7. Performance of different methods benchmarked using DM nsSNVs from HGMD and rare variants from gnomAD.**

### Supplementary Note

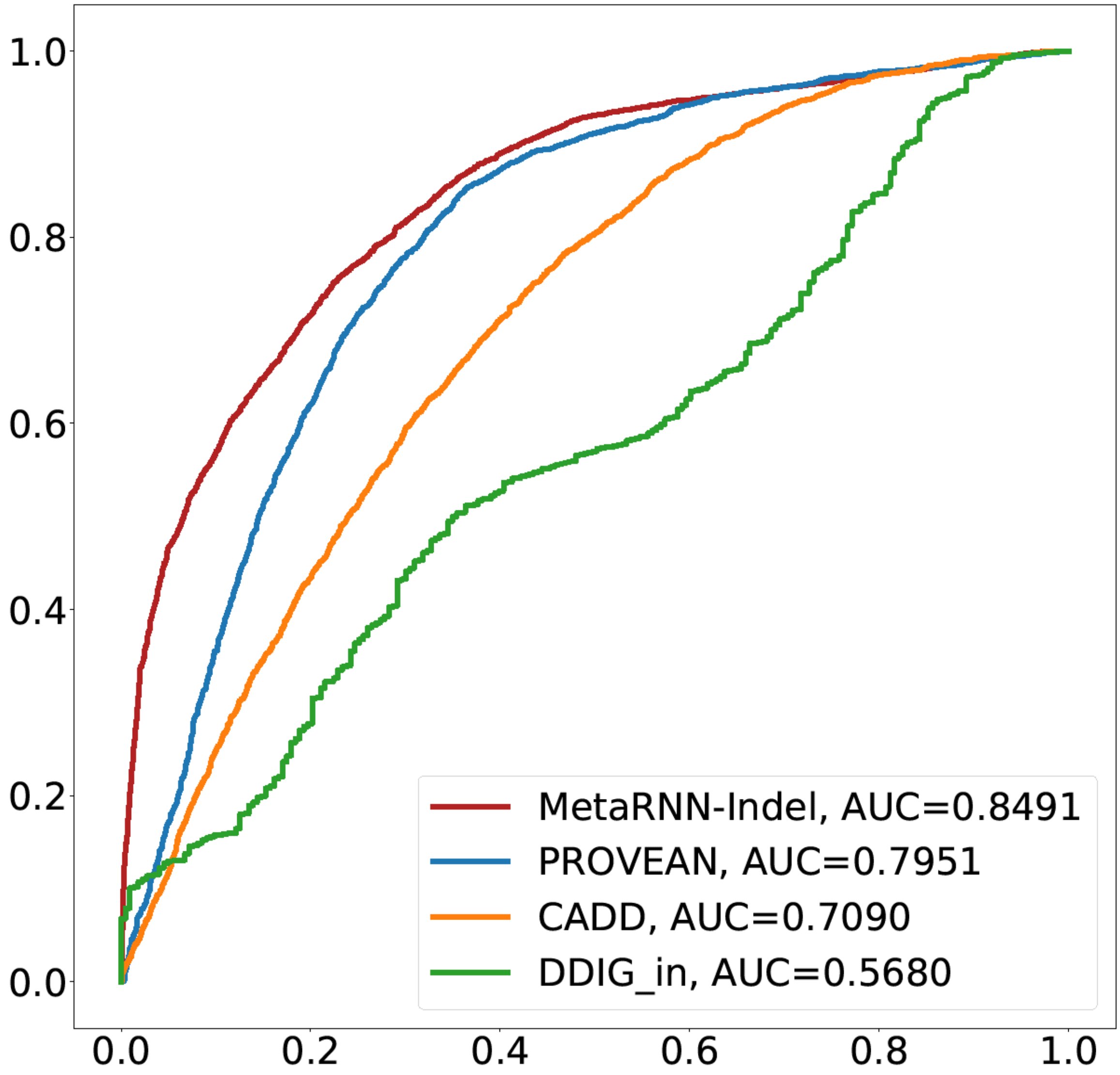


**S. Figure 8. Performance of different methods benchmarked using DM nfINDELs from HGMD and rare variants from gnomAD.**

#### Supplementary Note

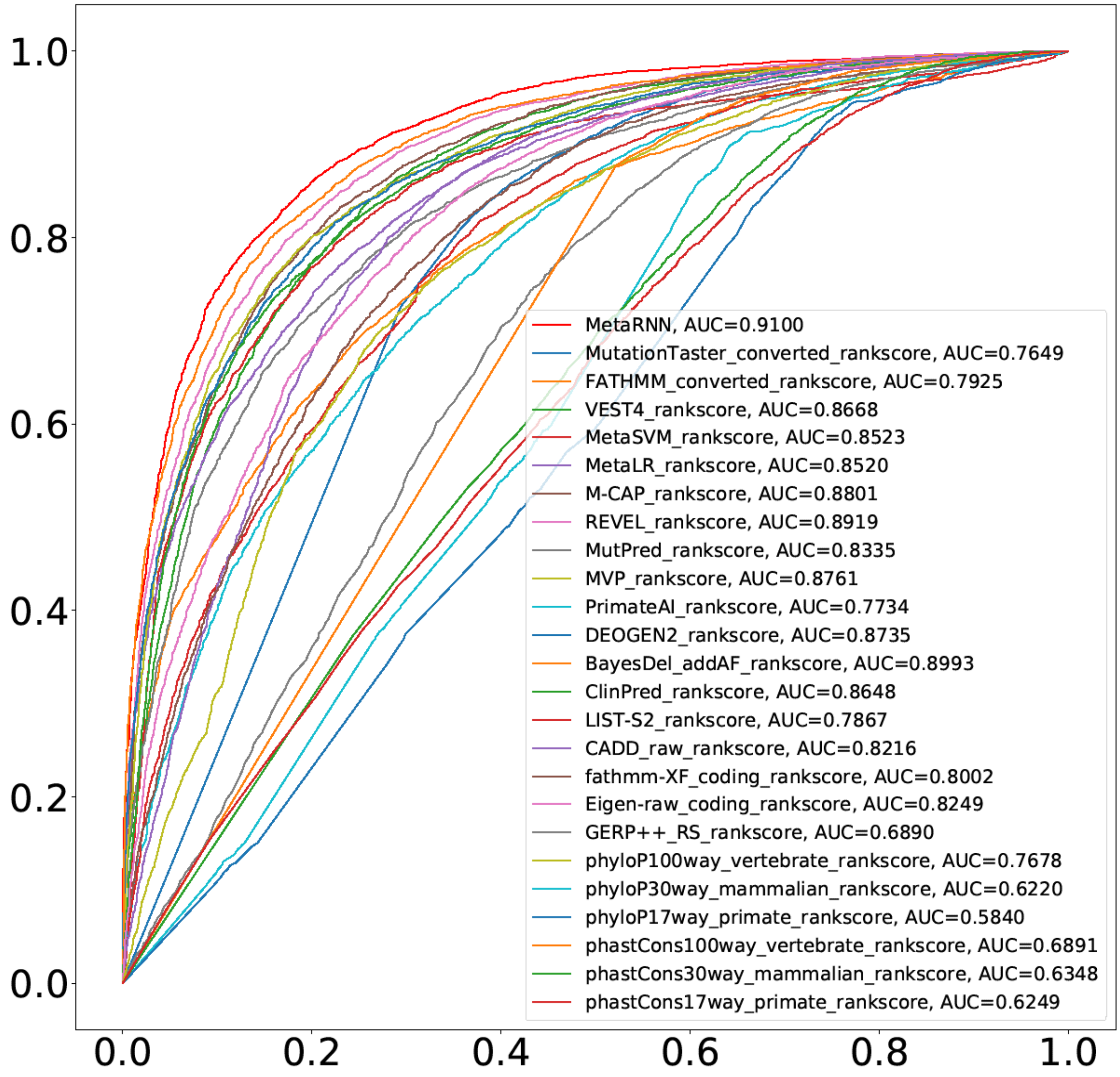


**S. Figure 9. Performance of different methods benchmarked using the rare nsSNV test set (RNTS).**

#### Supplementary Note

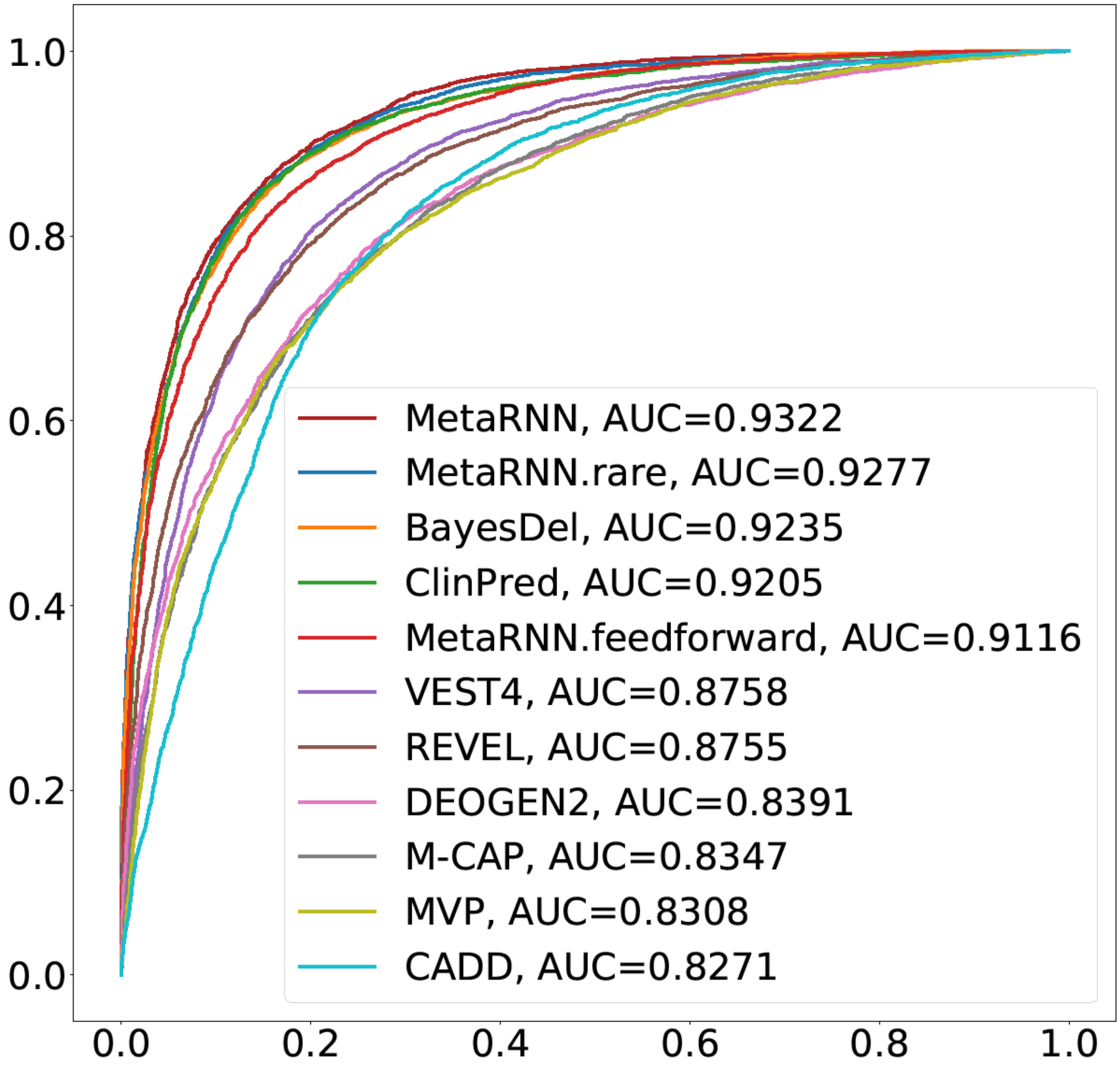


**S. Figure 10. Performance of different model structures and selected predictors benchmarked using the allele-frequency-filtered rare nsSNV test set (AF-RNTS).**
